# Liquid Crystalline Ordering of Banana-Shaped Gapped DNA Duplexes

**DOI:** 10.1101/2025.11.17.688902

**Authors:** Sineth G. Kodikara, Prabesh Gyawali, James T. Gleeson, Antal Jakli, Samuel Sprunt, Hamza Balci

## Abstract

In-phase adenine-tracts (A-tracts) introduce intrinsic bending to double stranded DNA resulting in banana-shaped macromolecules. In this study, we investigate how such sequence-dependent bending influences DNA-based liquid crystalline (LC) phases formed by gapped DNA (GDNA) constructs. By incorporating three in-phase A-tracts into each duplex arm, we created a GDNA construct with bent duplexes and examined the LC phases they form using temperature-resolved synchrotron small-angle X-ray scattering and polarizing optical microscopy. Like the analogous constructs containing straight (rod-like) duplexes, the bent constructs exhibit a transition from bi-layer smectic-B phase to a monolayer smectic-A phase, although at ~30 °C lower temperatures. By comparing the monolayer spacing between the bent and straight constructs, we estimate a bending angle of ~11° per A-tract at room temperature and at physiologically relevant salt and c_DNA_. The bending angle decreases with increasing DNA concentration and temperature. Moreover, we demonstrate that divalent cations enhance the stability of the smectic-B phase up to ~30 mM Mg^2+^ but reduce it beyond that. The reduced thermal stabilities of the bilayer and in-layer ordering of bent duplexes imply reduced propensity for DNA condensation and heterochromatin formation under physiological conditions.

## Introduction

The material characteristics of double-stranded DNA (dsDNA) are closely intertwined with its physiological functions. Certain structural features have well-defined roles, such as the inward-facing bases of the double helix that shield the genetic code, while others are intriguing. For instance, despite its relatively large persistence length, dsDNA is highly bendable, allowing it to wrap around histone proteins. Despite its negatively charged backbone, dsDNA can undergo condensation (1) or form liquid crystalline (LC) phases under physiologically relevant DNA concentration (c_DNA_), temperature, ionic strength, and pH (2–4). Self-assembly and end-to-end stacking of short DNA fragments, and the spontaneous orientational ordering of the resulting rod-like aggregates creates a protective environment that stabilizes the otherwise fragile DNA molecules. These concepts might have played a role in the origin of life (5, 6).

A particularly underexplored structure–function relationship in dsDNA is *sequence-induced* bending of the duplex and its physiological implications. Certain sequence patterns introduce local curvature in dsDNA. For example, poly(A)-poly(T) base pairs (“A-tracts”) confer intrinsic curvature and increased rigidity to the duplex, which plays a key role in chromatin organization, including nucleosome depletion at these sites (7–10). A-tracts with a certain length and arrangement promote DNA condensation and enhance side-by-side attraction between duplexes in the presence of cations (11–13). These modified interactions also significantly alter the LC behavior of DNA molecules that contain sufficiently long A-tracts compared to those with mixed sequences (14).

When arranged ‘in-phase’ with the helical pitch (such as A_x_N_10-x_A_x_ where N can be T, G, or C), A-tracts produce curved, banana-shaped constructs (15, 16) that resemble “bent-core” LC materials (17, 18). An example demonstrating the utility of this structural feature is the formation of DNA mini-circles (~10 nm radius) used to confine proteins or lipids (19, 20). The bending observed in in-phase A-tracts has been attributed to narrowing of the minor grooves, arising from negative roll and large propeller twist angles between adjacent base pairs within A-tracts (21–23). The propeller twist angles describe rotation of two bases within one base pair relative to each other and are ~15-20° within A-tracts rather than the typical 10° (distinct from helical twist angles of 34-36° in the B-form DNA). These large propeller twist angles are associated with the formation of bifurcated, cross-strand hydrogen bonds, which in turn stabilize the highly twisted geometry. The uniformity of AT stacking within A-tracts allows for this conformation to persist coherently over several base pairs. This geometry tilts the base pairs toward the minor groove, effectively narrowing it. A consequence of the narrowed minor groove is allowing formation of a structured hydration layer (spine of water) in which water molecules bridge bases and phosphates across the groove. To summarize, the bifurcated hydrogen bonds and the structured hydration of the minor groove help stabilize the bending in A-tracts which occurs both within the A-tracts and at their interface with flanking sequences (24, 25). Extensive literature documents A-tract bending using experimental techniques such as NMR (26), scanning force microscopy (27), gel electrophoresis (28), fluorescence polarization anisotropy (29), single-molecule FRET (30), and computational approaches (31). These studies estimate a bending of 10°–18° per A-tract and report a reduction in bending angle with increasing salt concentration (28).

In this paper, we investigate the effect of duplex bending on the LC phase behavior in concentrated solutions of gapped DNA (GDNA) constructs (32), formed by linking two rigid dsDNA arms with a flexible single-stranded spacer (‘gap’). Figure 1A shows the construct designed for our study. Two 48 bp duplexes are connected by a 10-nucleotide long single strand (10 thymines). To introduce bending in the duplex segments, we incorporated an A_6_N_4_A_6_N_4_A_6_ sequence pattern (where N = T, G, or C), which is in-phase with the helical pitch, into the central region of the duplex arms. The detailed sequences of the DNA strands are given in the Supporting Information. We refer to the construct as b48-10T-b48 (“b” standing for bent).

**Figure 1:**
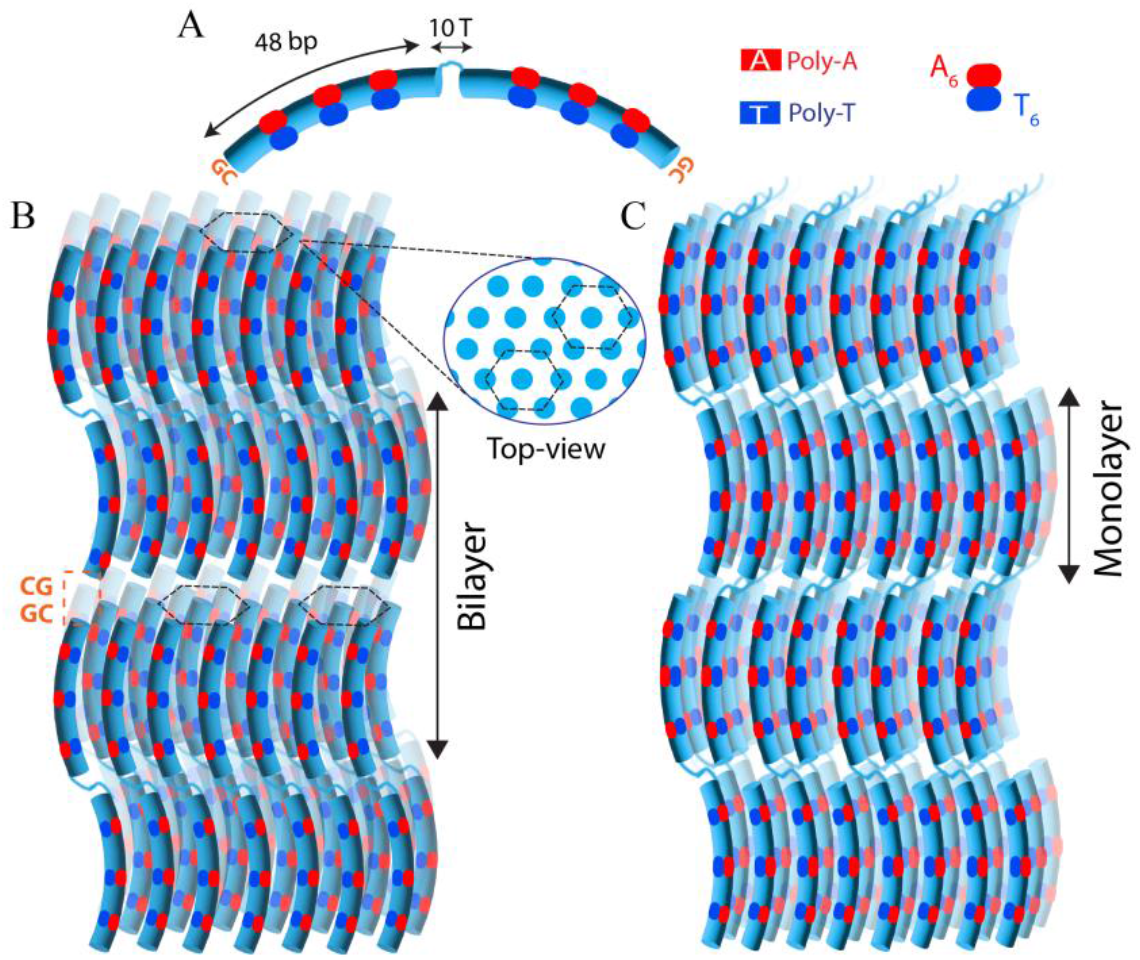
Schematic motifs of b48-10T-b48 GDNA constructs, which have 48 bp long bent duplex arms connected with a flexible 10T long spacer (“gap”). Each duplex arm has three in-phase A-tracts (A_6_N_4_A_6_N_4_A_6_N_4_) at its center which results in the bent configuration. Schematics showing (B) bilayer; and (C) monolayer smectic phases. The inset in (B) shows a schematic of the in-plane positional order (hexagonal). The semi-transparent molecules show the additional molecules within the layers in the back. The schematic in (B) shows the bilayer smectic phase for GDNA in the unfolded configuration, while Supporting Figure S1 shows the corresponding schematic for the folded configuration.

In previous studies on LC phase formation in solutions of 48-10T-48 GDNA with straight, rod-like duplex segments (mixed sequence), we observed a monolayer smectic-A phase at a DNA concentration (c_DNA_) of 260 mg/mL (33) and a bilayer smectic-B phase at c_DNA_ = 285 mg/mL (34). Both phases were observed at ambient temperatures and in the presence of monovalent cations (150 mM NaCl). The bilayer stacking has a periodicity approximately twice the length of a duplex segment (Figure 1B and Supporting Figure S1), while the monolayer has a periodicity of approximately the length of single duplex segment (Figure 1C). In the smectic-B phase the duplexes pack laterally within the layers in a hexagonal lattice, while in the smectic-A phase the layers are liquid-like (short range positional correlations within the layers).

In the current study, we investigate smectic ordering in b48-10T-b48 solutions with c_DNA_ in the 265–295 mg/mL range using synchrotron small-angle X-ray scattering (SAXS) and polarizing optical microscopy (POM). Additionally, we study the effect of divalent cations – specifically, the introduction of 0–100 mM MgCl_2_ – on the smectic ordering of b48-10T-b48 GDNA. We demonstrate that GDNA constructs with bent duplexes form smectic phases similar to those of GDNA with rod-like duplexes (33–35); although, with significantly reduced stability. We also report measurements for the bending angle per A-tract as a function of temperature, in the 5–50 °C range, and of c_DNA_. The physiologically relevant c_DNA_ examined in this study is typically inaccessible using alternative approaches. Finally, we demonstrate that despite the duplex bending, Mg^2+^ ions effectively stabilize both the in-layer positional order and end-to-end stacking interactions similarly to our previous finding in dense solutions of “straight” 48-10T-48 GDNA (36).

## Results and Discussion

Figure 1A shows a schematic of the GDNA constructs where the bending introduced by the in-phase A-tracts is assumed to be distributed throughout the duplex structure. Figure 2A-D show temperature dependent SAXS data for the b48-10T-b48 construct at 150 mM monovalent salt (NaCl) concentration and for DNA concentrations c_DNA_=265, 275, 285 and 295 mg/mL. The data covers the temperature range 5–65 °C. At T=5 °C and c_DNA_=275 mg/mL, two prominent peaks are observed at *q*_1_ ≈ 0.192 ± 0.003 nm^−1^ and *q*_2_ ≈ 0.396 ± 0.003 nm^−1^. While the peak amplitudes vary significantly with c_DNA_, their positions remain largely unchanged and correspond to layer spacings of 32.7 ± 0.5 nm and 15. 9 ± 0.1 nm, respectively. This suggests while the mass density contrast (which determines the peak amplitude) in the direction perpendicular to the planes has a strong dependence on c_DNA_, the interlayer spacing does not. Given that the length of a 48 bp mixed sequence duplex is about 16 nm, we designate the peaks at positions *q*_1_ and *q*_2_ as bilayer and monolayer peaks, respectively (Figure 1B-C). The peak at *q*_2_ represents an overlap (within resolution of our experiment) of the second order peak due to bilayer phase and the first order peak of the monolayer phase. Figure 3 shows POM images that exhibit azimuthally striated fan textures at T=20 °C. The fan texture is also present at 40 °C and 52 °C, but the striation pattern is washed out at these higher temperatures. These fan textures are consistent with a smectic layer structure (34, 37).

**Figure 2:**
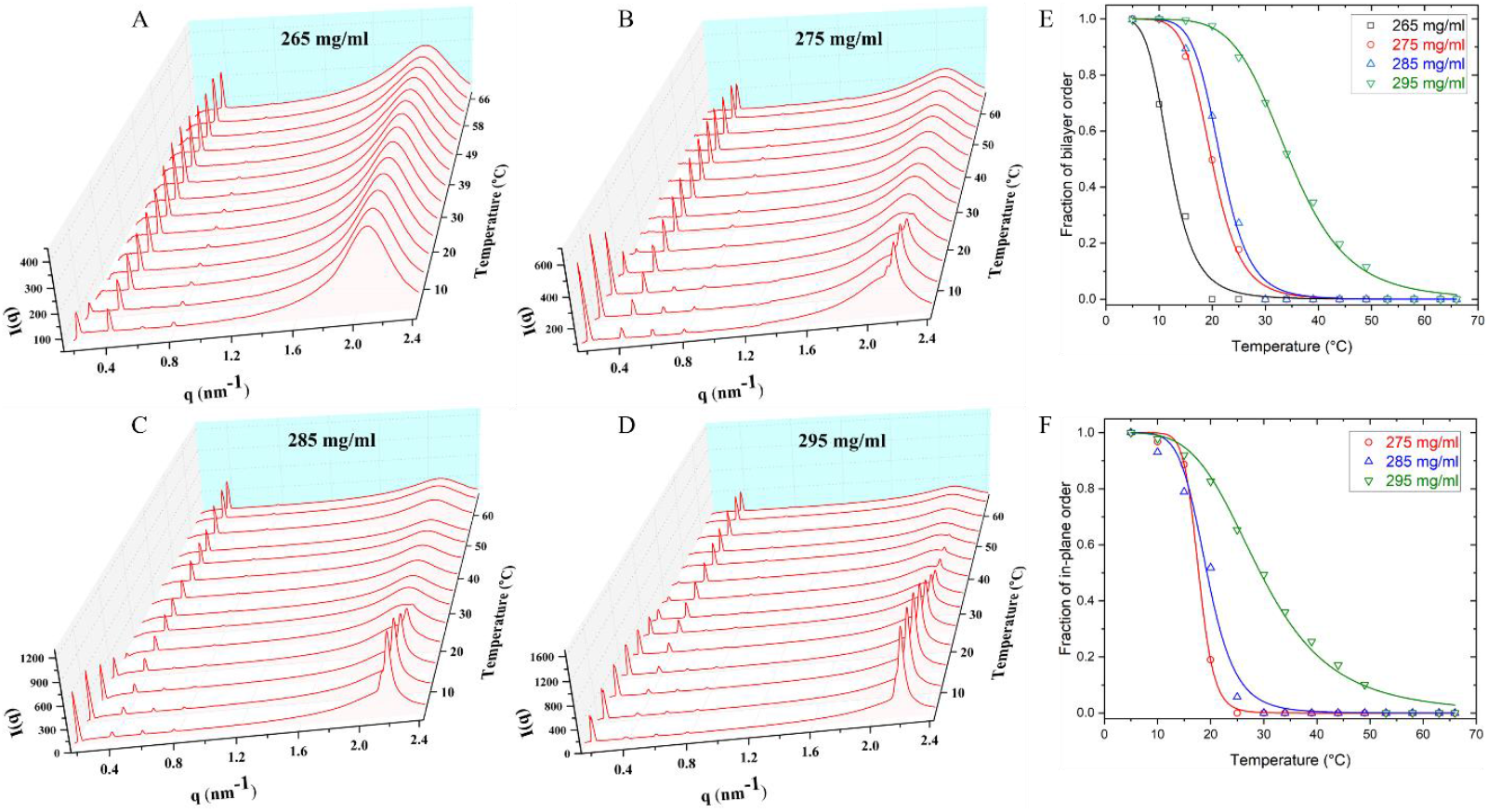
Temperature dependence of the azimuthally averaged SAXS intensity vs. scattering wavenumber *q* for b48-10T-b48 samples with varying c_DNA_. The data were acquired on a heating cycle (gradually increasing temperatures) at c_DNA_ of (A) 265 mg/mL; (B) 275 mg/mL; (C) 285 mg/mL; and (D) 295 mg/mL. All samples (except c_DNA_=265 mg/mL) are in a bilayer smectic-B phase at low temperatures. This phase melts at elevated temperature, as indicated by diminishment of small-angle peak at *q*_1_ *≈* 0.19 *nm*^−1^ (that is specifically associated with the bilayer spacing) and by the disappearance of the wide-angle peak at *q*_*w*_ *≈* 2.2 *nm*^−1^ (which is associated with in-layer positional ordering of the duplexes). The c_DNA_=265 mg/mL construct shows a bilayer phase but no in-plane order at T≈7 °C. (E) Fraction of bi-layer phase as a function of temperature and associated Hill fit to determine the thermal melting temperature for samples with different c_DNA_. (F) Similar analysis for the fraction of in-plane order vs. temperature. The thermal stability of the bilayer stacking and in-plane order increases with increasing c_DNA_. Details of the fitting analysis are provided in Supporting Information and the fit results are reported in Table 1.

**Figure 3:**
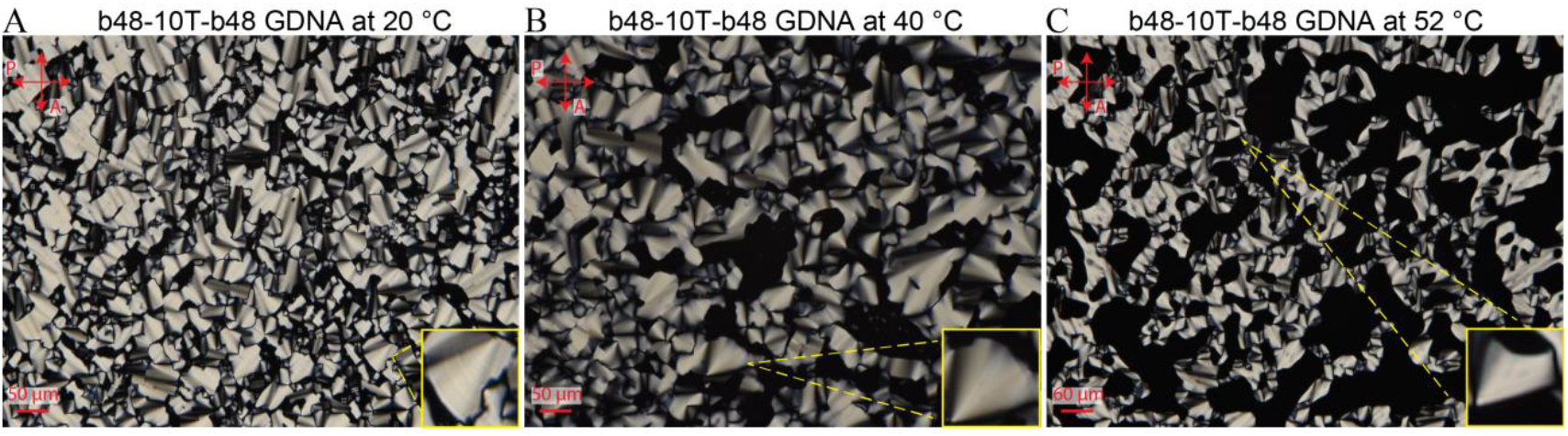
POM images of a b48-10T-b48 sample in 150 mM NaCl solution at (A) 20 °C; (B) 40 °C; and (C) 52 °C. The insets highlight a magnified view of the fan texture, which is characteristic of smectic phase. The azimuthal striation that is present at 20 °C is washed out at 40 °C and 52 °C data. These experiments were repeated at least 5 times, and several hundred images were acquired at different regions.

Consistently with 48-10T-48 and 48-20T-48 constructs (14, 34), the bilayer structure and in-layer positional order associated with the smectic-B phase for the b48-10T-b48 construct become more prominent and stable as c_DNA_ increases (Figure 2A-D). For the 48-10T-48 constructs, the smectic-B phase was observed for c_DNA_=285 mg/mL (34) at *T* ≈ 5 °C but not at c_DNA_=260 mg/mL (33). Similarly, for b48-10T-b48, the bilayer order develops at *T* ≤ 10 °C and c_DNA_=265 mg/mL (Figure 2A) and persists up to *T* ≤ 40 °C for c_DNA_=295 mg/mL (Figure 2D). The smectic-B phase is replaced with a monolayer smectic-A phase with layer spacing slightly less than a single duplex segment at elevated temperatures. The observed stabilities of the positional order are significantly lower in bent constructs compared to straight constructs. This suggests (i) weaker end-to-end stacking interactions, likely resulting from imperfect alignment of duplex ends; (ii) less dense in-plane packing, as indicated by increased helix–helix separation (smaller *q*_*w*_); and (iii) stretching of the 2D lattice along one axis, as evidenced by the splitting of the wide-angle peak (discussed below).

**Table 1.** Results for the characteristic melting temperature *T*_*m*_ from the fits using a single Hill function and described in the Supporting Information.

|  |  | Bilayer | In-plane |
| --- | --- | --- | --- |
| $c_{\text{DNA}}$ (mg/mL) | [MgCl<sub>2</sub>] (mM) | $T_m$ (°C) | $T_m$ (°C) |
| 265 | 0 | 12.0±0.2 | N/A |
| 275 | 0 | 19.9±0.2 | 17.8±0.1 |
| 285 | 0 | 21.6±0.3 | 19.3±0.4 |
|  | 10 | 66.7±1.7 | 61.7±1.4 |
|  | 30 | 66.7±2.9 | 69.2±1.8 |
|  | 100 | 19.8±0.2 | 18.3±0.4 |
| 295 | 0 | 34.5±0.3 | 29.4±0.5 |

While the temperature-resolved SAXS experiments can be analyzed to determine the impact of bent duplexes on LC phase stability, a comparison of the monolayer spacings of the bent and straight constructs can be used to quantify the bending induced by the in-phase A-tracts. For this, we compared the monolayer spacing exhibited by b48-10T-b48 with that of 48-10T-48 solutions (that terminate with AT base pairing in both duplex arms) at the same c_DNA_. SAXS spectra for the latter are presented in Supporting Figure S2. We modeled the bending of b48-10T-b48 duplexes as a circular arc with a continuous bend (Figure 4A). Assuming each A-tract contributes equally to the total bending, θ = 3θ_*b*_, where θ_*b*_ is the bending angle per A-tract. In this model, the contour length (*ℓ*) of the duplex arm is *ℓ* = *r*(2θ). The contour length represents the monolayer spacing for the straight, rod-like duplex. Similarly, the end-to-end separation is chord of the arc: *c* = 2*r* sin(θ), which represents the monolayer spacing for the bent duplex. The ratio of the two monolayer spacings, 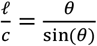, can be solved numerically to determine the bending angle per A-tract θ_*b*_. The result of this analysis is presented in Figure 4B where variation of bending angle per A-tract is plotted as a function of temperature for different c_DNA_ conditions. The temperature is restricted to the range in which the bilayer phase is stable since the second harmonic peak of this phase overlaps with the first harmonic peak of the monolayer phase (2*q*_1_ = *q*_2_). Beyond this temperature, a sharp decrease is observed in the calculated bending angle, which we believe is an artefact to the melting of the bilayer phase (Supporting Figure S3). The uncertainties in θ_*b*_ were calculated from the uncertainty in peak position corresponding to the spacing between *q* values recorded on the SAXS detector. The bending angle decreases with increasing c_DNA_ and temperature. The partial loss of narrowing of the minor grooves and disruption of the structured water spine that stabilizes the A-tract geometry have been proposed as the main drivers for the reduced bending at higher temperatures (21, 38).

**Figure 4:**
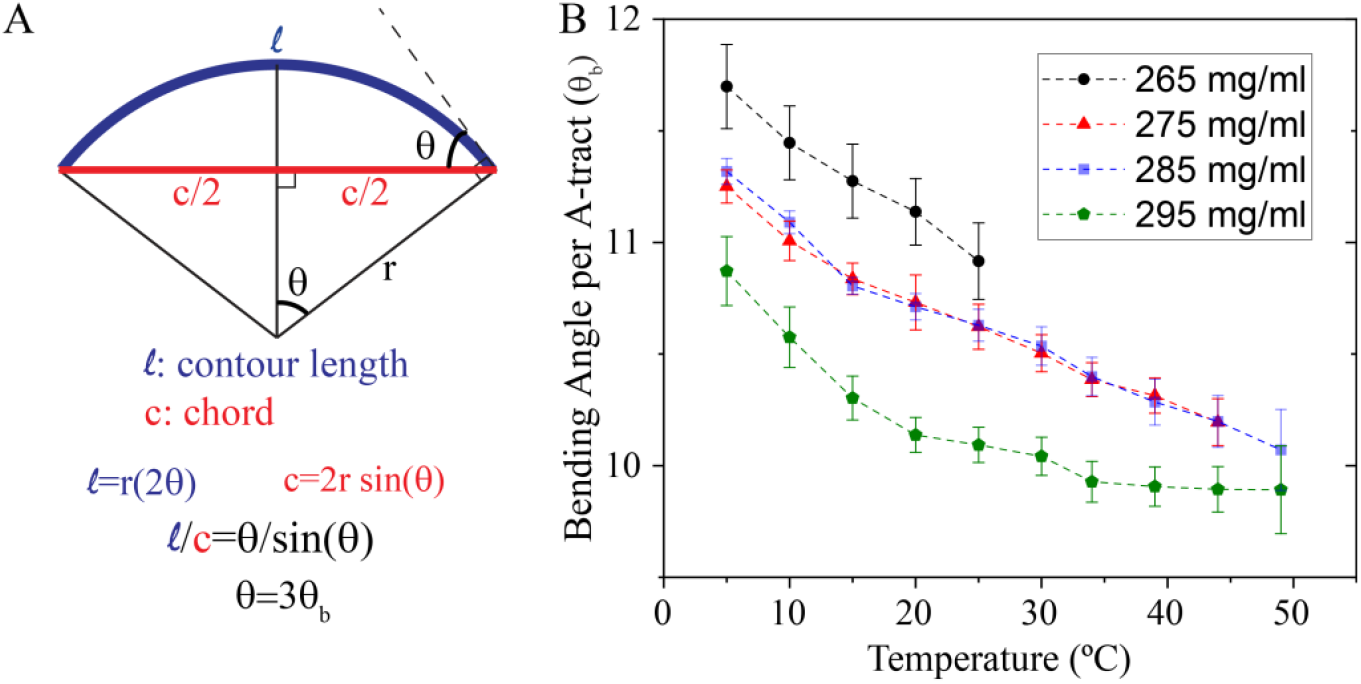
Bending angle calculations. (A) The bending in the duplex arms that contain three in-phase A-tracts is modeled as a continuous bend (circular arc) with a total bending angle of θ, where θ=3θ_*b*_ (assuming equal bending of θ_*b*_in each A-tract). The contour length of the black thick line represents the length of straight duplex, which is determined based on the monolayer spacing of GDNA with rod-like duplexes. The thick red line is the chord and represents the end-to-end separation of a bent duplex which is determined based on the monolayer spacing of the GDNA with bent duplex arms. (B) Bending angle per A-tract θ_*b*_ is numerically calculated assuming the model described in (A). The data for all c_DNA_ are shown until the melting of their respective bilayer phase. The bending angle decreases with increasing temperature, by about ~10% in the 5-50 °C range for c_DNA_=275-295 mg/mL.

The bending angle of θ_*b*_ ≈ 11° at room temperature is at the lower end of the range (10-18°) typically reported in the literature. However, given the decrease in bending with increasing c_DNA_ and the significantly greater c_DNA_ used in our measurements compared to those accessed in other methods, a lower value is to be expected. Notably, our bending angle estimate applies for physiologically relevant *c*_*DNA*_ and is therefore relevant to crowded (cellular) environments. Supporting Figure S4 shows an alternative model in which the bent duplex is modeled to have 3-kinks located at the center of each A-tract, which results in larger angles.

The wide-angle diffraction peak in Figure 2 signifies in-layer positional ordering of the bent duplexes in b48-10T-b48 solutions. Its amplitude and position at *q*_*w*_ ≈ 2.15 nm^−1^, are similar to the peak associated with hexagonal positional ordering previously observed in 48-10T-48 solutions at the same c_DNA_ (34). Assuming a hexagonal packing, the peak in b48-10T-b48 solutions corresponds to an average lateral duplex-duplex separation of *a* = 4*π/*√3*q*_*w*_ ≈ 3.37 *nm*, quite comparable to the 3.29 *nm* lateral spacing between 48-10T-48 constructs (28). There is, however, a significant difference in the wide angle scattering. Closer examination of the peak reveals a splitting of the wide angle peak into two narrowly separated peaks at *q*_*w*1_ ≈ 2.13 nm^−1^ (*a* = 3.41 *nm*) and *q*_*w*2_ ≈ 2.18 nm^−1^ (*a* = 3.33 *nm*) at T=5 °C. Supporting Figure S5 shows an example of this for a b48-10T-b48 solution with c_DNA_=285 mg/mL. The splitting, which was not observed in 48-10T-48 GDNA with straight duplexes, can be explained by the packing scheme shown in Figure 1B, where the duplex arms of the b48-10T-b48 constructs all bend in the same plane perpendicular to the layers. This would produce a slight stretching of the 2D lattice along the axis parallel to this plane and to the layers, breaking the sixfold symmetry and resulting in basis vectors of different lengths.

To quantify the thermal stability of the smectic-B phase, we determined the associated melting temperatures (*T*_*m*_, listed in Table 1) for the bilayer stacking and the in-layer positional ordering using a procedure described previously (35) and outlined in the Supporting Information. The corresponding results of this procedure are shown in Figure 2E-F. The in-layer positional order generally melts at slightly lower temperatures than the bilayer stacking, which is consistent with previous findings for GDNA with straight duplexes (35, 36). To assess the impact of bending, we can compare the *T*_*m*_ values for b48-10T-b48 with those of 48-10T-48. Although the temperature-dependent SAXS spectra of 48-10T-48 constructs with AT terminal base pairs were published in an earlier study (34), their thermal stability was not quantified; this is now presented in Supporting Figure S6. A comparison of the *T*_*m*_ values shows that the bending causes a significant reduction (at least 30 °C) in the thermal stability of the smectic-B phase. For instance, at c_DNA_=285 mg/mL, the melting temperature of the bilayer stacking is *T*_*m*_ = 52.5 ± 0.2 °C for 48-10T-48 compared to a significantly lower value *T*_*m*_ = 21.6 ± 0.3 °C for b48-10T-b48. Similarly, the respective melting temperatures for in-plane positional order are *T*_*m*_ = 48.9 ± 1.6 °C and *T*_*m*_ = 19.3 ± 0.4 °C (Supporting Figure S6). The pronounced decrease in the stability of the bilayer smectic phase suggests that end-to-end stacking interactions are substantially less effective due to the increased curvature of the duplexes, likely leading to imperfect alignment of their blunt ends. Furthermore, the slightly larger lateral duplex–duplex separation and the disruption of six-fold symmetry (as indicated by the splitting of the *q*_*w*_ peak) imply that in-plane packing is also less efficient in the bent constructs, further contributing to the reduced stability. These variations would be further amplified if the bent and rod-like duplexes had consistent terminal base pairs: our b48-10T-b48 constructs have GC-GC terminal base pairs, which enhance the stability of end-to-end stacking (35) relative to the AT-AT terminals on the 48-10T-48 construct.

An interesting trend in the b48-10T-b48 constructs is the absence of biphasic melting behavior, which was observed in GDNA constructs with straight duplexes and required a double Hill function with two temperatures *T*_*m*,1_ and *T*_*m*,1_ to account for the shape of the melting curves (35). This was attributed to stacking interactions in different configurations, i.e., the terminal XY base pairs of duplexes in each half of a bilayer could stack in XY:XY or XY:YX configurations. With bent duplexes, this heterogeneity is largely eliminated, possibly because only GC:CG stacking orientation is stable enough to maintain the bilayer smectic phase.

Previously, we reported on 48-20T-48 constructs containing 10 consecutive poly(A)-poly(T) sequences at the center of each duplex arm, which are not in-phase with the DNA helix and therefore do not induce significant bending (14). These constructs exhibited a novel columnar phase at lower temperatures (*T* < 35 °C) and a remarkably stable bi-layer smectic-B phase at higher temperatures (*T* > 35 °C). This sharp contrast with the phase behavior of b48-10T-b48 solutions at comparable c_DNA_ has a straightforward explanation: When the b48-10T-b48 constructs are efficiently packed to form a 2D hexagonal lattice with periodicity comparable to the duplex diameter, layering in the third direction is simultaneously imposed, and columnar order is sterically frustrated.

The ionic content of the environment is another key factor influencing both DNA condensation and the stability of LC phases. Figures 5A-D show temperature-resolved SAXS spectra for the b48-10T-b48 construct at c_DNA_=285 mg/mL with 0, 10, 30, and 100 mM MgCl_2_, in addition to a constant 150 mM NaCl. Figures 5E-F present the analysis for determining the thermal stability (with *T*_*m*_ values listed in Table 1). The thermal stability of both the bilayer stacking and in-plane positional order increases with MgCl_2_ concentration between 0-30 mM but decreases back to the 0 mM level at 100 mM MgCl_2_. This trend is consistent with previous observations for GDNA constructs with straight duplexes, suggesting that the distribution of divalent cations along the dsDNA backbone follows a similar pattern, despite the bending induced by A-tracts.

**Figure 5:**
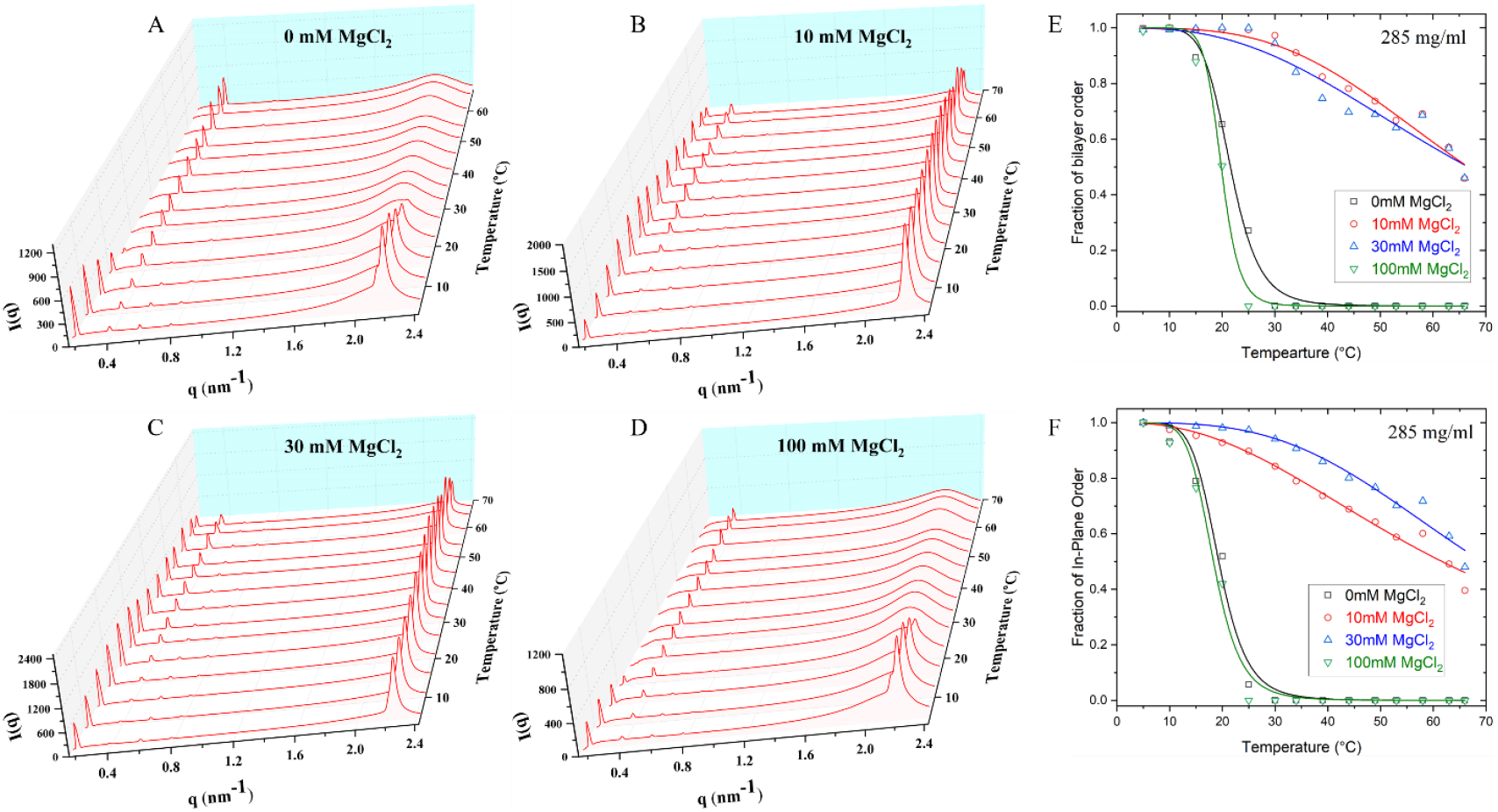
Temperature dependence of the azimuthally averaged SAXS intensity vs. scattering wavenumber *q* for b48-10T-b48 samples at c_DNA_=285 mg/mL and with varying MgCl_2_ concentration. The data were acquired on a heating cycle at MgCl_2_ concentration of (A) 0 mM; (B) 10 mM; (C) 30 mM; and (D) 100 mM. All samples exhibit a bilayer smectic-B phase at low temperatures, which is replaced with a monolayer phase at higher temperatures. (E) Fraction of bilayer domains as a function of temperature and associated Hill fit (see Supporting Information) used to determine the melting temperature for samples with different c_DNA_. (F) Similar analysis for the fraction of in-plane order vs temperature. The thermal stability of the bi-layer phase and in-plane order increases with MgCl_2_ concentration up to 30 mM but significantly decreases at 100 mM. The fit results are reported in Table 1.

Such a non-monotonic dependence on divalent cation concentration has been observed in all-atom molecular dynamics (MD) simulations investigating the condensation of short duplexes (12, 13). These MD simulations proposed DNA–ion–DNA bridging models in which multivalent cations that localize (or are adsorbed) on one DNA molecule establish attractive interactions with the negative charges on the phosphate backbone of another molecule (13, 39). In this model, spatial correlations between the multivalent cations that localize on the major grooves of DNA are prevalent at lower cation concentrations, and result in effective attractive interactions, which we interpret to result in elevated stability of LC phases. These simulations show that such correlations are washed out at higher concentrations and result in a uniform positive charge distribution that manifests as repulsive interactions between DNA molecules. We propose the nonmonotonic dependence on Mg^2+^ concentration observed in our experiments to be a result of a similar mechanism.

While the Mg^2+^ ions are primarily expected to shield the helix-helix repulsion within the smectic planes, it is worth noting the remarkable increase in the stability (a shift of over 40 °C in *T*_*m*_) of both in-plane and bi-layer smectic ordering with the introduction of 10 mM MgCl_2_, which attests to a strong coupling of the helix-helix and end-to-end stacking interactions. How the bending angle is impacted by Mg^2+^ ions is an important question that would need further studies where the monolayer spacing for 48-10T-48 constructs is measured as a function of MgCl_2_ concentration. Another interesting feature of the 10 and 30 mM MgCl_2_ data is the absence of a clear splitting of the wide-angle peak within the resolution of our diffraction measurements. Evidently, the additional shielding provided by Mg^2+^ ions reduces the anisotropy in packing of bent duplexes within the layers.

While not investigated in this study, the length of the flexible single stranded spacer (‘gap’) connecting the duplex arms is a determining factor in the formation and stability of smectic phases formed by GDNA. We explored this issues earlier (32) and showed that (for the investigated c_DNA_ range) the spacer length should be at least 10 nt for formation of bilayer smectic phase and at least 4 nt for formation of monolayer smectic phase. We project the minimum spacer length (for a given c_DNA_) that is required for formation of smectic phases would be greater for GDNA with bent duplexes. The lower stability induced by stacking and helix-helix interactions in the bent-duplexes would require a greater entropic effect. This could be provided by compartmentalization of a longer flexible spacer into a volume in which it can attain greater conformational entropy.

## Conclusion

Our study leads to several key conclusions. First, we demonstrate that SAXS studies on LC phases formed by GDNA can resolve temperature-dependent variation of A-tract induced DNA bending under physiologically relevant ion and DNA concentrations. The bending (~11° per A-tract at T=25 °C) is at the lower end of the range typically reported in the literature and decreases with increasing c_DNA_ and temperature. The physiologically relevant crowding achieved in our studies makes these measurements better estimates for the level of bending induced by in-phase A-tracts in the cellular setting. Second, the cumulative bending (~33°) introduced in the duplex arms results in significant reduction in the stability of LC phases and variations in the in-plane packing compared to the case of GDNA with straight duplexes. These suggest such sequences could impact biomolecular organization in a physiological setting. Finally, introducing modest concentrations of Mg^2+^ results in a remarkable increase in the stability of both in-plane order and bilayer stacking in the smectic-B phase, highlighting the significance of helix-helix interactions in these condensed phases of DNA. With our growing know-how about these systems, we envision making alterations to the GDNA constructs or the environmental conditions that would modulate these interactions in a predictable manner. Our study illustrates the exceptional sensitivity of LC phases formed by GDNA to structural and environmental variations, further cementing its potential as a versatile tool to study related concepts in DNA condensation and biomolecular organization.

## Acknowledgements

The research reported here was supported by the National Science Foundation under grant DMR-1904167. The authors are particularly grateful to Ruipeng Li and Masa Fukuto for their assistance in performing the SAXS/WAXS measurements on the CMS beamline (11-BM) at the National Synchrotron Light Source II, a U.S. Department of Energy (DOE) Office of Science User Facility operated for the DOE Office of Science by Brookhaven National Laboratory under Contract No. DE-SC0012704.

## Supporting Information Available

Details of GDNA synthesis and various data analysis methods, sequences of DNA constructs, and SAXS measurements on 48-10T-48 construct, an alternative method to estimate the bending angles, data on splitting of the wide-angle peak

## Notes

### Competing Interest Statement

The authors have declared no competing interest.

