## Supporting Information for "Liquid Crystalline Ordering of Banana-Shaped Gapped DNA Duplexes"

### Materials And Methods

**Gapped DNA (GDNA) Synthesis:** DNA oligomers (O1, O2, O3, O1b, O2b, and O3b) were purchased from Genescript (Piscataway, NJ) in polyacrylamide gel electrophoresis (PAGE) purified form. The sequences of all oligos are given in the DNA Sequences section below.

GDNA synthesis and sample loading into thin-walled borosilicate capillaries were carried out as described previously (Gyawali et al., 2021). The GDNA constructs (b48-10T-b48) consist of two symmetric 48-base pair duplexes connected by a 10 nt long single-stranded DNA gap of consecutive thymine bases. To create the bent GDNA constructs, a long strand (O1b, 106 nt) is annealed with two short strands (O2b and O3b, each 48 nt) which are complementary to either side of O1b except the 10T sequence in the middle. Similarly, the GDNA with straight, rodlike duplexes was created by annealing O1, O2, and O3. After annealing, the GDNA solutions are passed through a 50 kDa membrane filter (Amicon Ultra from Millipore) to remove incomplete constructs and diluted such that they contain ~30 mM NaCl by adding distilled and deionized water. This solution is then concentrated by centrifuging it for 12 min at 12,000 rpm (GDNA concentration increases while NaCl concentration is kept at ~30 mM). The GDNA concentration is measured with a NanodropOne Spectrometer (Thermo Fisher Scientific) and is typically in the 80-100 mg/ml range at this step. This solution is then added to quartz capillaries (2 mm inner diameter) with an open end. Water is slowly evaporated which results in increasing GDNA and NaCl concentration. The NaCl concentration is adjusted such that ~150 mM NaCl will be reached when the desired DNA concentration is achieved.

**SAXS Measurements:** SAXS measurements were carried out on beamline 11-BM at the National Synchrotron Light Source II. The incident X-ray energy was 17 keV, and the incident beam size at the sample was 0.2 × 0.2 mm. The typical acquisition time for SAXS patterns was 5 s, which did not result in visible X-ray damage to samples. A commercial hot/cold stage with Kapton film windows was used to regulate the sample temperature between 5-65 °C, which is well below the thermal melting temperature (~78 °C at the relevant DNA concentration) of the 48-bp duplex arms. We also recorded background scattering from a capillary containing pure buffer solution, which was subtracted from the data taken on the GDNA samples during processing of the SAXS patterns. A capillary filled with silver behenate powder was used to calibrate the scattering wave number ($q$) in the plane of the detector.

**Polarizing Optical Microscopy:** Polarizing optical microscopy was performed on $\sim 5-10$ $\mu$m thick samples sandwiched between microscope slides. The samples were sealed by a thin ring of mineral oil after water had evaporated from the edges of the film to the point where focal conic (FC) textures appeared, signaling the development of a smectic phase.

**DNA Sequences:** The b48-10T-b48 GDNA constructs with bent duplexes were formed by annealing the following three oligos of 106 nt (O1b) and 48 nt (O2b and O3b):

O1b: GCAGATGCACATAAAAAAGTGGAAAAAACTTGAAAAAAGTACTTCGAA TTTTTTTTTT AAGCTTCATGAAAAAAGTTCAAAAAAGGTGAAAAAATACACGTAGACG

O2b: TTCGAAGTACTTTTTTCAAGTTTTTTCCACTTTTTTATGTGCATCTGC

O3b: CGTCTACGTGTATTTTTTCACCTTTTTTGAACTTTTTTCATGAAGCTT

The 48-10T-48 GDNA constructs with straight, rodlike duplexes were formed by annealing the following three oligos of 106 nt (O1) and 48 nt (O2 and O3):

O1: ACAGATGCACATATCGAGGTGGACATCACTTACGCTGAGTACTTCGAA TTTTTTTTTT AAGCTTCATGAGTCGCATTCACTACAGGTGGAGCTATACACGTAGACA

O2: TTCGAAGTACTCAGCGTAAGTGATGTCCACCTCGATATGTGCATCTGT

O3: TGTCTACGTGTATAGCTCCACCTGTAGTGAATGCGACTCATGAAGCTT

**Sequence Variations Among Bent and Straight Duplexes:** For clarity, we have color coded the sequence variations between the bent and straight duplexes using O1b and O1 strands below. Green sequences are common while the red/black are the variations.

O1b: G*CAGATGCACAT*-AAAAAA-*GTGG*-AAAAAA-*CTT*G-AAAAAA-*GTACTTCGAA*-*TTTTTTTTTT*-*AAGCTTCATG*-AAAAAA-G*TTC*-AAAAAA-*GGTG*-AAAAAA-TACACGTAGACG

O1: A*CAGATGCACAT*-ATCGAG-*GTGG*-ACATCA-*CTT*A-CGCTGA-*GTACTTCGAA*-*TTTTTTTTTT*-*AAGCTTCATG*-AGTCGC-A*TTC*-ACTACA-*GGTG*-GAGCTA-*TACACGTAGAC*A

As illustrated, there is a variation of four nucleotides in this 106 nt long construct other than the poly-A repeats, which are required in order to induce significant bending in the duplexes. So, we consider the poly-A repeats as a required and essential modification for studying LC phases formed by bent GDNA and determining the bending angles.

Two of the four nucleotide modifications were made in the immediate vicinity of A-tracts (A to G, underlined) to avoid breaking the $A_6N_4A_6N_4A_6$ pattern. Not changing these A's to G's would have resulted in an $A_6N_4A_6N_3A_7$ pattern in both duplex arms. So, we consider this also to be a necessary modification.

The remaining two nucleotide variation are two terminal bases: G…G in the bent constructs vs. A…A in the straight constructs. The G…G terminal bases result in the more stable GC-GC terminal base stacking interactions while A…A results in the significantly less stable AT-AT terminal base stacking. Anticipating a reduced stability due to bending of the duplexes, we opted for GC-GC termination to increase the probability of bilayer smectic phase and in-plane positional order formation over an extended temperature range. The stability of LC phases of bent constructs with AT-AT termination is expected to be much lower (as much as 30 °C in the GDNA constructs with straight duplexes) and it is not clear whether they would form over a meaningful temperature range to enable the type of analysis performed in this study. In the particular case of bending angle calculation, the terminal base pairs are not expected to influence the results since the bending angles are calculated based on monolayer spacing (rather than stability), which has not shown a measurable variation with changes in terminal bases.

**Thermal Melting Analysis:** Thermal melting analysis is performed as described in more detail previously (Kodikara et al., 2023). This analysis relies on the thermal evolution of the area under the relevant SAXS peaks, which are the first (fundamental) and third order peaks for the bi-layer order and the sharp wide-angle peak for the in-layer order. The cumulative area under the first and third order peaks were used to characterize the bi-layer phase. The square root of the area scales with the magnitude of the density modulation or order parameter that characterizes the degree of positional order in each case. Each peak was fit to a Gaussian function in $q$ using Origin software, and the total area under the Gaussian was then calculated. The square root of the resulting total integrated intensity under the relevant peaks is normalized with respect to the relevant maximum value and then plotted as a function of temperature. The results were fit to single and double Hill functions of the form,

$f(T) = 1 - \frac{T^n}{T_m^n + T^n}$ and $f(T) = 1 - \frac{1}{2}\left(\frac{T^n}{T_{m1}^n + T^n} + \frac{T^k}{T_{m2}^k + T^k}\right)$ , where $f(T)$ is the average fraction of "bound" duplexes, $T_m$, $T_{m1}$, and $T_{m2}$ are characteristic melting temperatures, and $n$ and $k$ are positive exponents. Table 1 lists these values for all the constructions we investigated. In certain cases, the double Hill function produces an artificial kink in a region where the data is sparse. With limited synchrotron beam time, it was not feasible to perform measurements at higher temperature resolution.

### Additional Schematics

Figure 1B shows a schematic of the bilayer smectic phase formed by a folded by GDNA, which consists of two 48-bp segments connected with a 10T linker (Figure S1A). A smectic phase with a similar bilayer separation can also be formed by unfolded GDNA in a different orientation (Figure S1B) or a GDNA in the folded conformation (Figure S1C). These different arrangements cannot be distinguished based on layer spacing alone.

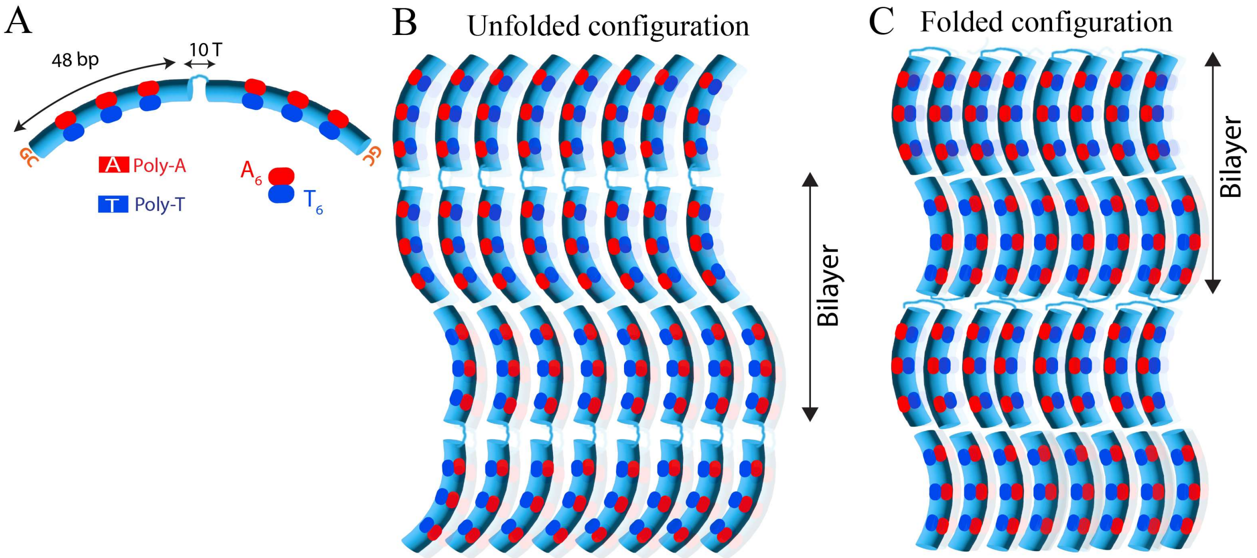


Figure S1. Schematics of (A) b48-10T-b48 GDNA construct. (B) bilayer phase in the unfolded configuration in which the bent in both duplexes within a GDNA are pointing in the same direction (they can also point in opposite directions as illustrated in Figure 1B). (C) bilayer phase in the folded conformation.

### SAXS Data of 48-10T-48 GDNA Constructs with AT-AT Terminal Base Pairs

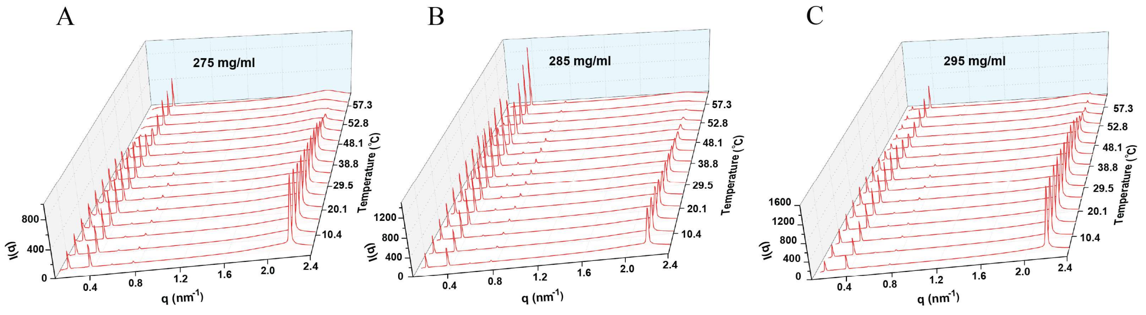


Figure S2. Temperature dependence of the azimuthally averaged SAXS intensity vs. scattering wavenumber *q* for 48-10T-48 samples with varying $c_{DNA}$. The data were acquired on a heating cycle (gradually increasing temperatures) at $c_{DNA}$ of (A) 275 mg/ml; (B) 285 mg/ml; (C) 295 mg/ml.

### Alternative Model for DNA Bending

We modeled the bending of b48-10T-b48 duplexes as three kinks of equal angles ($\theta_b$) at the center of the A-tracts, i.e., located 14, 24, and 34 bp away from either duplex end (Figure S3A). In this model, the contour length ($\ell$) of the duplex arm is $\ell = 4.8x$ where $x = 10\ bp$. The contour length represents the monolayer spacing for the straight, rod-like duplex. Similarly, the end-to-end separation ($c$) is $c = 1.4x\cos(3\theta_b/2) + x\cos(\theta_b/2) + x\cos(\theta_b/2) + 1.4x\cos(3\theta_b/2)$, which represents the monolayer spacing for the bent duplex. The ratio of the two monolayer spacings, $\frac{\ell}{c} = \frac{2.4}{1.4\cos(3\theta_b/2)+\cos(\theta_b/2)}$, can be solved numerically to determine the bending angle per A-tract $\theta_b$. Figure S3B shows the results of this analysis, which yields the same trends as those of circular arc model presented in Figure 4. The bending angle per A-tract is $\theta_b \approx 16°$ at room temperature. Similar to Figure 4, the bending angles are shown until the melting of their respective bilayer phase.

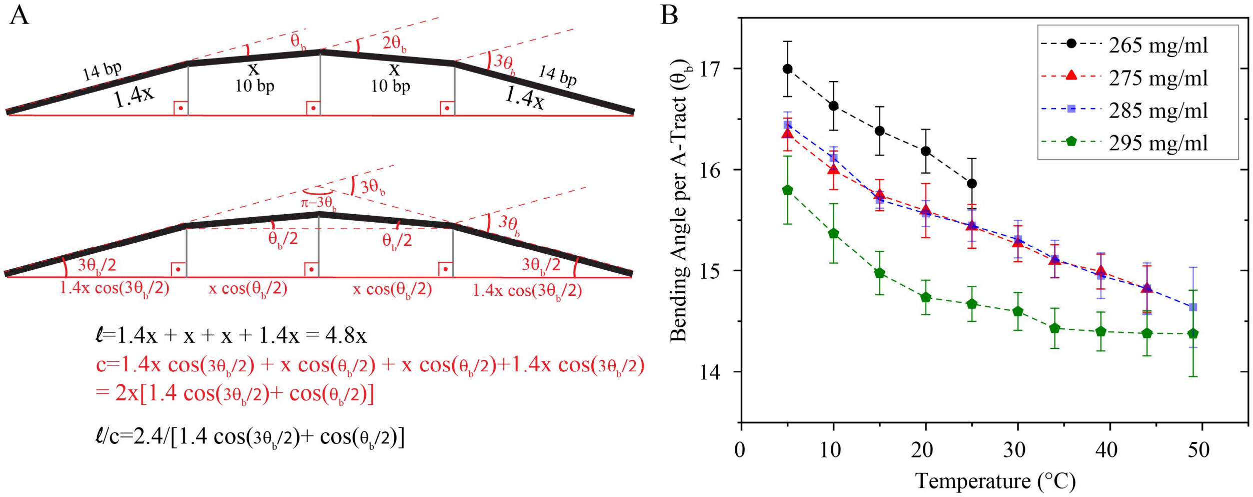


Figure S3. Bending angle calculations. (A) The bending in the duplex arms that contain three in-phase A-tracts is modeled as three kinks of equal bending ($\theta_b$) localized at the center of the A-tracts. The contour length of the black thick line represents the length of straight duplex, which is determined based on the monolayer spacing of GDNA with straight, rod-like duplexes. The thick red line represents the end-to-end separation of a bent rod that includes three kinks. The length of the red line is determined based on the monolayer spacing of the GDNA with bent duplex arms. (B) Bending angle per A-tract $\theta_b$ is numerically calculated assuming the model described in (A). The data for all $c_{DNA}$ are shown until the bilayer phase melts.

### Impact of Smectic-B Phase Melting on Bending Angle

Fig. S4A shows the bending angle $\theta_b$ as a function of temperature for the circular arc model for an extended temperature range that shows a sharp drop in the bending angle during melting of the bilayer phase. This sharp drop in $\theta_b$ is an artifact of our inability to track the position of the fundamental peak for the monolayer phase ($q_2$, which is used in calculating the bending angles) at the temperature range of bilayer phase melting. Fig. S4B-D illustrate the coincidence of the melting of the bilayer peak with the sharp drop in the bending angle for $c_{DNA}$=275 mg/ml, 285 mg/ml, and 295 mg/ml, respectively. The second order of the bilayer phase ($2q_1$) overlaps with the first order peak of the monolayer phase ($q_2 \approx 2q_1$). Melting of the bilayer phase results in a rapid decrease in the amplitude of the $2q_1$ peak over a relatively narrow temperature range (~50-65 °C). As the two peaks are not resolved, we cannot confidently establish the position of $q_2$ peak. Given these constraints, our method of determining the bending angle should not be relied upon in the 50-65 °C range in which the bilayer phase melts. This temperature has also been avoided in most experimental studies measuring the bending angle, likely because it is high enough to result in transient melting of the base pairs. While this may not be critical for the LC phases, it is important for bending angle calculations which strongly depend on base pairing interactions and their specific geometry.

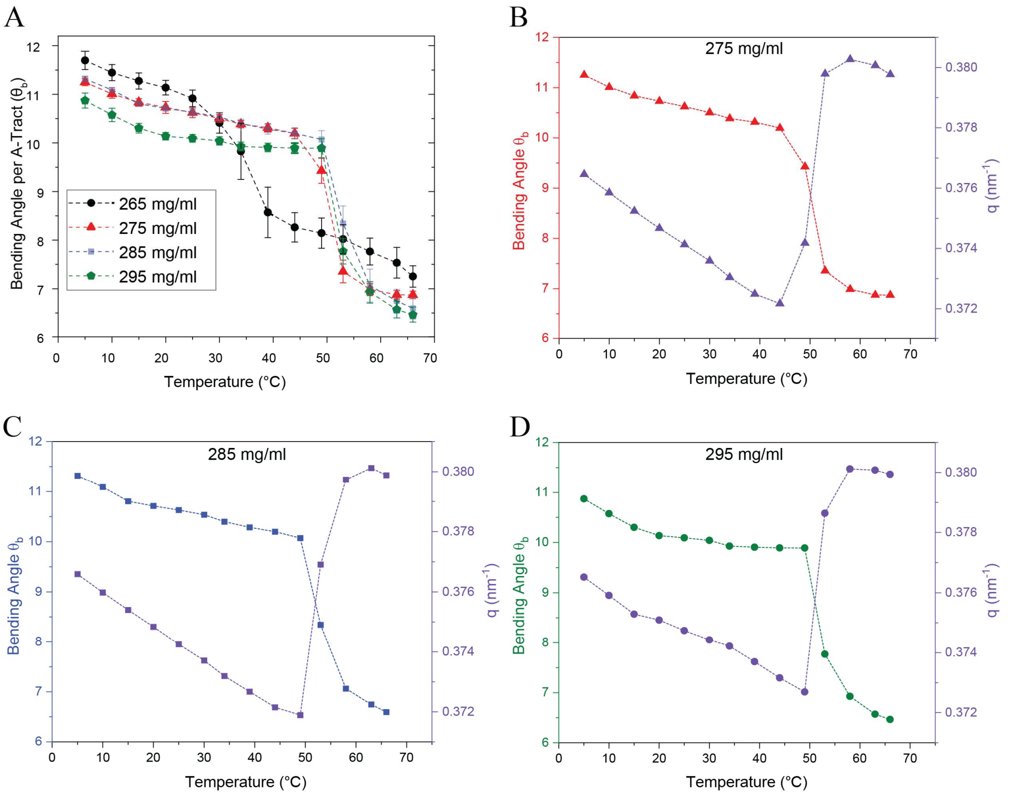


Figure S4. (A) Bending angles for the circular arc model are shown over an extended temperature range that includes melting of the bilayer phase (a narrower range is shown in Fig. 4B). The sharp decrease in $\theta_b$ at T $\approx$ 50 °C for $c_{DNA}$=275-295 mg/mL is due to the shift in the monolayer peak position (which is used for calculating $\theta_b$) when the overlapping first harmonic of the bilayer smectic phase melts. (B-D) Data showing the shift in wavenumber (q) of the monolayer peak position when the bilayer smectic-B phase melts at different DNA concentrations: (B) 275 mg/ml, (C) 285 mg/ml, and (D) 295 mg/ml. The shift in monolayer peak position to higher $q$ values in the bent construct (which takes place at higher temperatures in the straight construct due to higher thermal stability) causes a sharp decrease in the bending angle.

### Splitting of the Wide-Angle Peak

The wide angle peak in the smectic-B phase of bent duplexes at $c_{DNA}$=285 mg/ml is split into two narrowly separated peaks at $q_{w1} \approx 2.13\ \mathrm{nm}^{-1}$ $(a = 3.41\ nm)$ and $q_{w2} \approx 2.18\ \mathrm{nm}^{-1}$ $(a = 3.33\ nm)$ at T=5 °C. Figure S5 shows evolution of these split peaks as the temperature is raised from 5 °C to 25 °C.

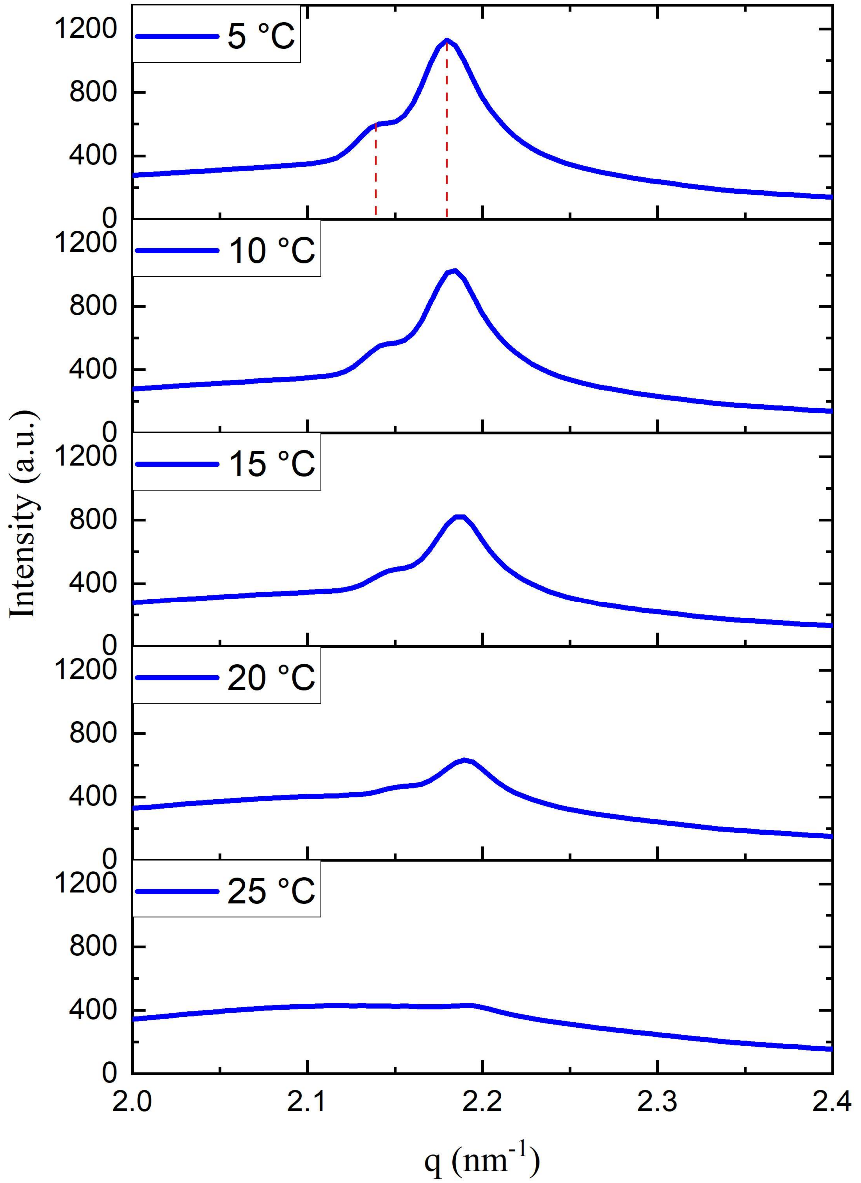


Figure S5. Evolution of the wide-angle peak splitting as the temperature is raised from 5 °C to 25 °C for the b48-10T-b48 sample at $c_{DNA}$=285 mg/ml (in the absence of $MgCl_2$).

**Thermal Melting Analysis of 48-10T-48 GDNA with AT-AT Terminal Base Pairs**

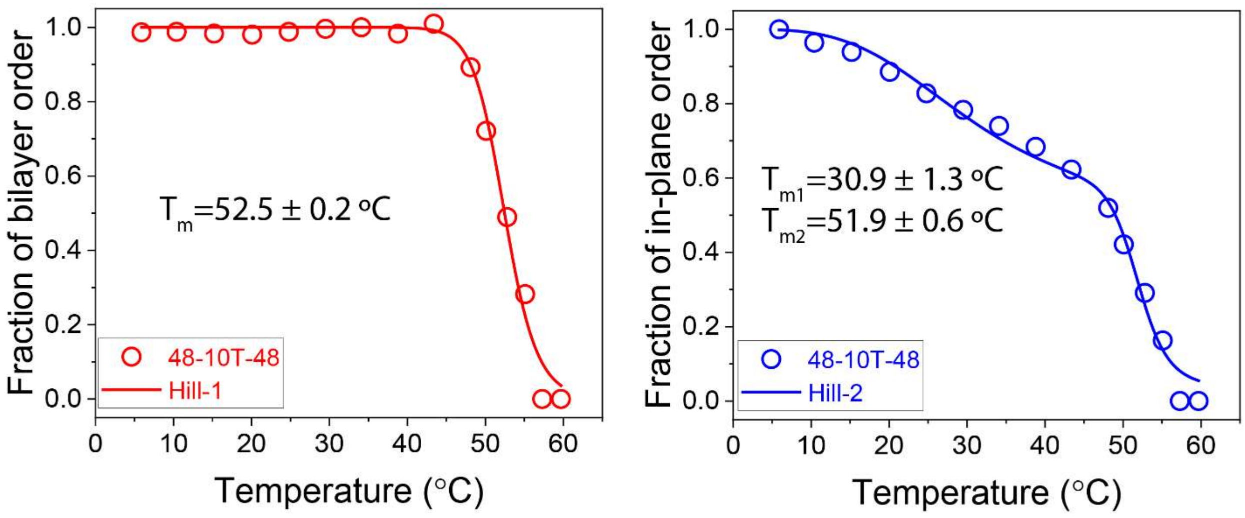


Figure S6. Thermal melting analysis for 48-10T-48 sample at 285 mg/ml. The associated SAXS data was published previously without the thermal melting analysis (Gyawali et al., 2022).
